# The VGNC: Expanding Standardized Vertebrate Gene Nomenclature

**DOI:** 10.1101/2022.08.10.503477

**Authors:** Tamsin E.M. Jones, Bethan Yates, Bryony Braschi, Kristian Gray, Susan Tweedie, Ruth L. Seal, Elspeth A. Bruford

## Abstract

The Vertebrate Gene Nomenclature Committee (VGNC) was established in 2016 as a sister project to the HGNC (HUGO Gene Nomenclature Committee), to approve gene nomenclature in vertebrate species without an existing dedicated nomenclature committee. The VGNC aims to harmonize gene nomenclature across selected vertebrate species in line with human gene nomenclature, with orthologs assigned the same nomenclature where possible. This article presents an overview of the VGNC project and discussion of key findings resulting from this work to date. VGNC approved nomenclature is accessible at https://vertebrate.genenames.org, and is additionally displayed by the NCBI, Ensembl and UniProt databases.

## Background

The HUGO Gene Nomenclature Committee (HGNC) (1) is the only organization worldwide that assigns standardized gene nomenclature to human genes, and has been doing so for over 40 years. HGNC has always coordinated closely with the other groups that are actively assigning standardized nomenclature to vertebrate model organisms, namely mouse (2), rat (3), chicken (4), Xenopus (5) and zebrafish (6). In all of these species the genes are named relative to their HGNC named human orthologs and paralogs, and in the vast majority of cases exactly the same gene symbols and names are adopted for orthologous genes. The main exceptions to this are genes within complex gene families where there have been multiple gene duplication and loss events throughout evolution, such as the olfactory receptors and the zinc fingers, where homology relationships can be difficult to establish without in-depth analysis. Note there are differences in capitalization to conform to each species’ conventions: mouse and rat gene symbols begin with an uppercase letter followed by lowercase letters, Xenopus and zebrafish symbols use lowercase letters only, and symbols in chicken, human and other mammals contain all uppercase letters.

In 2004, to facilitate a project to improve the links between orthologous human and mouse genes, HGNC created the HGNC Comparison of Orthology Predictions (HCOP) tool (7). The HCOP tool aggregates predicted orthologs to human genes from a number of expert orthology resources that use differing methodologies. The consensus output from HCOP enabled HGNC curators to link orthologous human and mouse genes and ensure they had the same gene symbols where possible. This tool was subsequently expanded to include further orthology resources and now includes a total of 20 species (8).

With the rapid release of a large number of genomes from well-studied vertebrates in the 2000s, it became clear there was a need for assigning standardized nomenclature in key species that were not being served by a dedicated nomenclature authority. In the absence of approved symbols, NCBI Gene and Ensembl routinely project gene nomenclature to predicted homologs but these are automated assignments that may not be based on approved or unique nomenclature, and may not be consistent between or even within resources. In October 2009 the HGNC organized the “Gene Nomenclature Across Species” meeting in Cambridge, UK, with invited participants from the fields of genome assembly and annotation, phylogenetics and gene naming (9). A key conclusion from this meeting was that a core set of consensus 1:1 orthologs between given species (especially human and another organism) should be derived through comparing data. This gene set could then be automatically named in line with the orthologs in an already named species. It was appreciated that this approach would not work for complex gene families, and so these would need to be named following expert manual curation.

The HGNC was clearly well placed to work on assigning names in selected species, with the HCOP tool and established links with both collaborating nomenclature groups in other species and with experts for specific complex gene groups. Following successful funding applications, this work commenced under the project name the “Vertebrate Gene Nomenclature Committee”. A pilot project was conducted with chimpanzee (*Pan troglodytes*) gene naming in the first instance, with every chimpanzee gene symbol and name undergoing manual review before approval in the VGNC database. Over 16,000 chimpanzee gene symbols have been approved to date. Six additional vertebrate species have since been added to the VGNC database: cattle (*Bos taurus*, added in 2017), horse (*Equus caballus*, added in 2017), dog (*Canis lupus familiaris*, added in 2017), cat (*Felis catus*, added in 2019), macaque (*Macaca mulatta*, added in 2019), and pig (*Sus scrofa*, added in 2020). The genes in these species are named via a combined approach of automated and manual approval via a curation tool. To date we have approved nomenclature for over 100,000 genes in these 7 species (see Table 1), as well as symbols for the complex cytochrome P450 gene family in a further 25 species (Additional file 1: Table S1).

**Table 1:** Number of genes approved in the seven core VGNC species as of January 2022.

| Species | # Approved Protein Coding Genes | # Approved Pseudogenes | # Approved Non Coding RNAs |
| --- | --- | --- | --- |
| *Pan troglodytes* | 16740 | 12 | 1 |
| *Bos taurus* | 16346 | 510 | - |
| *Equus caballus* | 15271 | 433 | - |
| *Canis lupus familiaris* | 15461 | 107 | - |
| *Felis catus* | 14017 | 2 | - |
| *Sus scrofa* | 13700 | 5 | - |
| *Macaca mulatta* | 14673 | 5 | - |

Gene nomenclature is only approved for genes that have gene models annotated in NCBI and/or Ensembl. An approved VGNC entry includes (but is not limited to) the approved gene name, approved gene symbol, a unique VGNC identifier in the format VGNC:##### where ##### is a number, the accession numbers for NCBI and/or Ensembl gene models, and links to orthologs in human and other species. Approved gene nomenclature is made publicly available on our website https://vertebrate.genenames.org (10), where users can choose to search and browse genes and gene groups in the form of “symbol reports” and “gene group reports” respectively, or alternatively download dataset files in their choice of file format. VGNC approved nomenclature is also disseminated by the NCBI, Ensembl and UniProt databases.

## Construction and Content

The process of nomenclature assignment is based on the identification of orthologs of human genes in the vertebrate species of interest followed by either automated nomenclature transfer or manual review by a curator. Orthology identification is performed using a subset of data from our HGNC Comparison of Orthology Predictions (HCOP) tool (8). The choice to start naming genes in a vertebrate species is informed by the quality of the genome assembly and annotation, its value as a model organism, and the level of interest from the community who work on the species. When a new species is added to the VGNC, a high confidence set of 1:1 orthologs with human is generated by comparing the results of four key orthology prediction resources in our HCOP tool: Ensembl Compara (11), NCBI Gene (12), Panther (13), and OMA (14). If all four resources agree on a 1:1 orthology relationship, the vertebrate ortholog is automatically assigned the human gene nomenclature, with some exceptions as outlined below. If three out of the four resources agree on a 1:1 ortholog, the vertebrate gene is marked for review by a curator before approval.

“Approving” a gene’s nomenclature in VGNC refers to a process whereby a gene is assigned an official gene symbol and name, allocated a unique VGNC ID, and made public on the VGNC website. Approving nomenclature for a gene also creates a link between the VGNC ID and the relevant NCBI and/or Ensembl gene models representing that gene, and results in the VGNC nomenclature being used for those gene models in their respective source databases.

We currently approve nomenclature in seven “core” VGNC species, that is, species for which we aim to assign nomenclature to the full protein-coding gene set. We also approve a subset of genes from specific complex gene families outside our seven core species when expertly curated data for these gene families are made available to us: at present this is limited to the cytochrome P450 genes, however we welcome gene family experts to contact us if they have curated gene family datasets across vertebrates that could be included in our database and therefore disseminated to other public databases.

VGNC data is stored in two separate databases, a production database that contains both approved VGNC genes and provisional VGNC genes (entries that require further evidence or curator input before they can be approved) and a release database that contains only those VGNC genes that have been approved and are publicly available on our website. An overview of the database schema is shown in Figure 1A. An automated analysis pipeline is run daily to update the data in these two databases and to ensure that they are synchronized both with each other and with any updates made to the nomenclature of orthologous human genes. The update pipeline is summarized in Figure 1B. Firstly, if any new genome assemblies are available for the core VGNC species from Ensembl or NCBI, the assembly information is added to an “assembly” database table. Chromosome and scaffold information is also imported into an internal curation tool (described below). Next, NCBI Gene data are updated to include any new, removed or modified gene entries for VGNC species, including location data. The same updates are made to Ensembl gene data if a new Ensembl release is available. Orthology data are imported via our HCOP tool and changes to existing data are updated. Changes to approved human gene nomenclature from HGNC are imported and used to automatically update orthologous VGNC gene entries in the VGNC database. Cross reference data such as UniProt IDs are updated and added to the relevant VGNC entries. Finally, the approved VGNC data are released to the public database and download files are updated on our FTP and Globus GridFTP servers.

**Figure 1:**
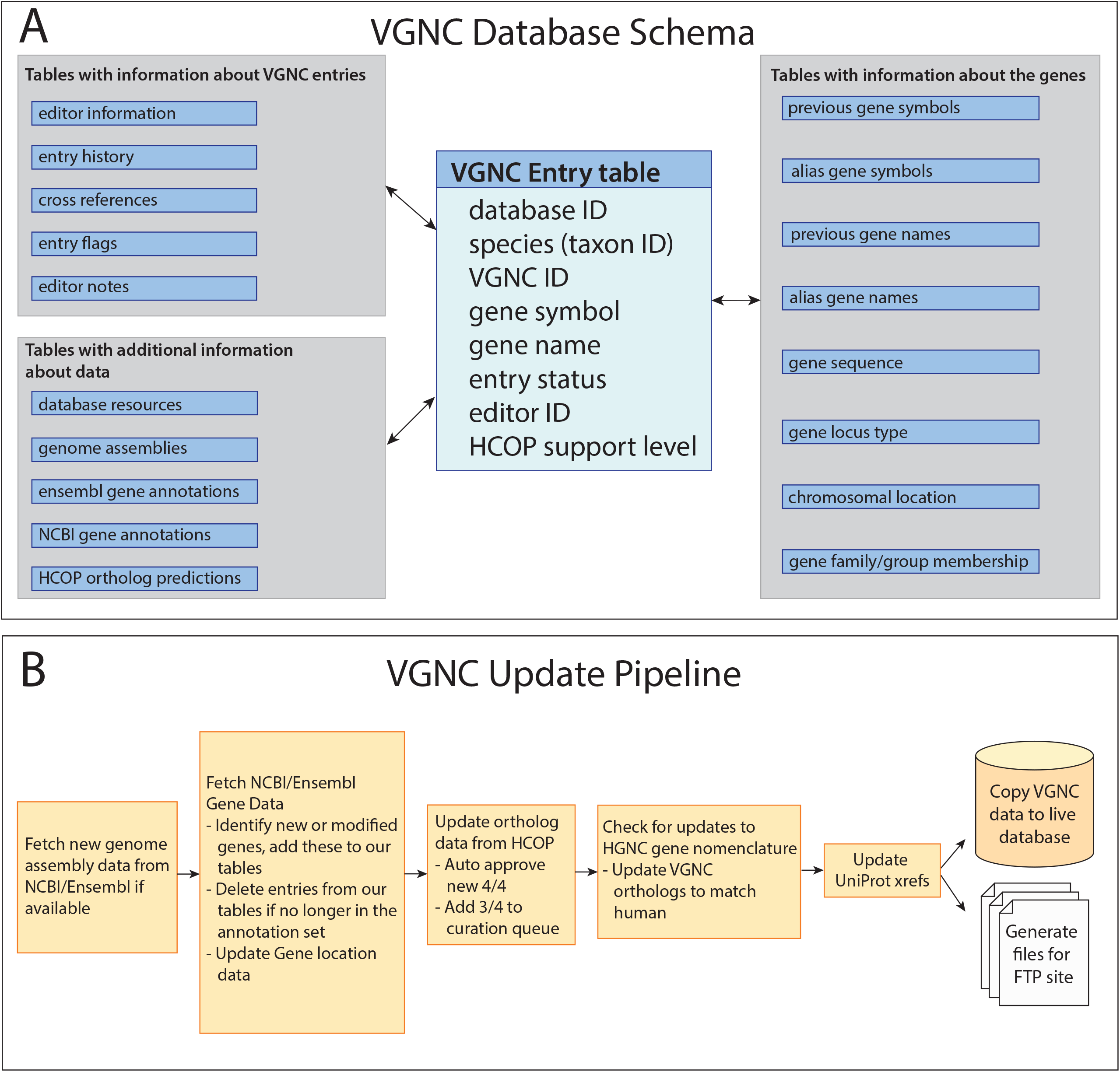
A: Simplified overview of the VGNC database schema. B: Overview of the VGNC update pipeline.

Not all human gene nomenclature is suitable for transfer to other species. We have already manually reviewed and updated many human gene names to make them species neutral but work on this is ongoing. Examples include genes that contain references to human disease or cancer: therefore, the VGNC pipeline implements a check for gene names containing terms like “syndrome” or “cancer” and prevents these from being automatically approved in VGNC. Instead these genes are added to the manual curation queue even where all four orthology resources agree on an orthology relationship.

We also prevent automatic approval of gene names referring to chromosomal location, eg “X-linked”, “Y-linked”, “region” or “neighbor”, since these may not apply in non-human species. If human gene nomenclature is updated in HGNC, the changes are usually automatically applied to any approved VGNC orthologs, unless a curator has manually marked the gene nomenclature to not be automatically updated. Examples of where this applies are where orthology is not 1:1 or where the human gene name contains a suffix of “(gene/pseudogene)” denoting it as a segregating pseudogene and this suffix should not automatically be transferred to non-human orthologs.

The process of manual curation is typically carried out using an internal curation tool that allows curators to compare the synteny and gene annotation models for a VGNC entry and its putative human ortholog as a starting point. Manual curation may also include conducting literature searches, consulting with experts, and phylogenetic analysis. If the gene annotations associated with a VGNC entry are syntenic with the human ortholog, the NCBI and/or Ensembl gene models appear to be accurate representations of the gene based on comparisons to orthologs, and the human gene nomenclature is suitable to transfer to other species, then a curator will manually approve the VGNC gene. If there are disruptions in synteny, further analysis is conducted to check if there may be deviations from a 1:1 orthology relationship. If there are issues with the NCBI and/or Ensembl gene models, such as locus type differences, merging of neighboring genes into a single gene model, or major discrepancies between the two models, it is sometimes possible to get errors corrected by contacting the relevant database; however, the VGNC entry will not be approved until there is at least one suitable gene model to approve as a cross reference. If there are issues with the human gene nomenclature, curators may consider modifying this nomenclature to make it transferable across species, or making a change to the VGNC gene nomenclature such that it differs from the human ortholog. For example, references to blood groups in human gene names are removed when transferring to non-human species. In the case of complex orthology relationships curators may perform phylogenetic analysis across several species and/or consult with experts before making nomenclature decisions for a given gene/gene family.

The VGNC collaborates with specialist researchers for certain complex gene families, particularly where orthology and paralogy relationships between species require careful study to discern. Two major examples are the Olfactory Receptors (15) and the Cytochrome P450s. Both of these gene families have undergone extensive manual curation in collaboration with expert advisors, and this work is ongoing. Briefly, expert advisors conducted comprehensive searches in each species for members of the relevant gene family and subsequently provided the VGNC with the genomic locations of all gene family members (including pseudogenes), and their suggested gene symbols for each gene. Each of these genes is manually confirmed by a VGNC curator, which can involve reviewing synteny comparisons, phylogenetic analysis, and identification of existing gene models in the NCBI and Ensembl genome annotation sets, or requesting the annotation of new gene models if none currently exist.

The VGNC coordinates with other gene nomenclature committees to approve consistent gene nomenclature across species where appropriate. This includes the Mouse Genome Nomenclature Committee (MGNC), the Rat Gene Nomenclature Committee (RGNC), the Chicken Gene Nomenclature Committee (CGNC), Xenbase, and the Zebrafish Nomenclature Committee (ZNC). Coordination across species is particularly crucial when naming genes in gene families where there is copy number variation and therefore 1:1 orthology relationships do not always exist. Examples of gene families whose nomenclature has been coordinated across all of the nomenclature committees include the oxytocin and arginine vasopressin ligand and receptor genes (16).

## Utility and Discussion

### Overview of VGNC Data

As of January 2022, there are 108,538 approved genes in VGNC. These include mostly protein coding genes (107,426) but we have approved a small number of pseudogenes (1,111) and 1 non-coding RNA. The non-coding RNA gene was originally approved as a protein coding gene but its locus type has since been updated to non-coding RNA. 107,283 genes have been approved in the seven core VGNC species. A summary of the number of genes approved per core species is shown in Table 1. A further breakdown of the numbers of automatically vs. manually approved protein coding genes in the core VGNC species is shown in Figure 2. VGNC also currently approves cytochrome P450 genes in a number of additional vertebrate species; the total number of genes approved in the 25 non-core species is 1,254 (see Additional file 1: Table S1 for a breakdown per species).

**Figure 2:**
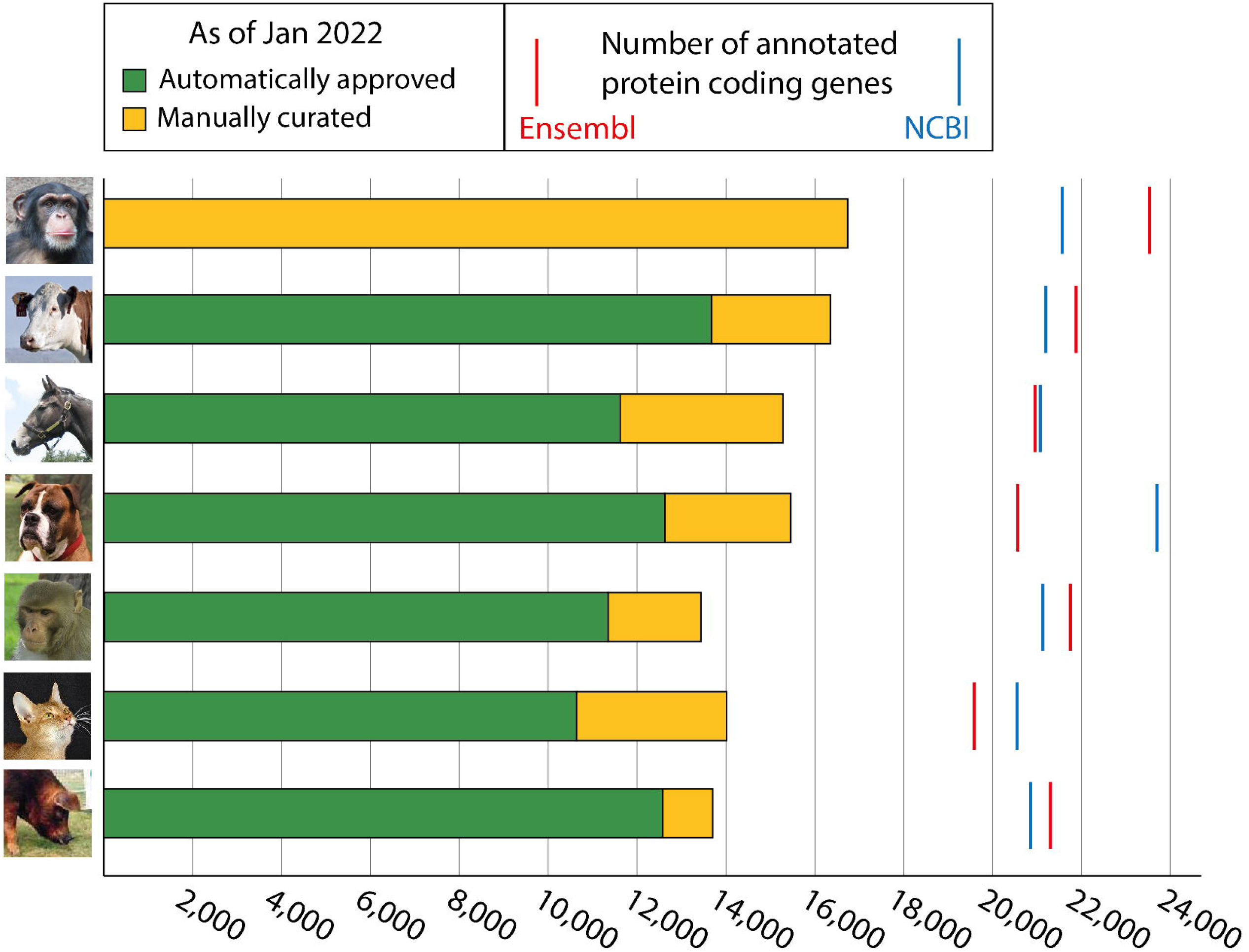
Numbers of automatically (green) and manually (yellow) approved protein coding genes in core VGNC species. Estimated total number of protein coding genes in the genome as annotated by Ensembl (red) and NCBI (blue) are indicated with lines. Based on genome assemblies: Chimpanzee - Pan_tro_3.0 (GCA_000001515.5, NCBI) Clint_PTRv2 (GCA_002880755.3, Ensembl); Cow - ARS-UCD1.2 (GCA_002263795.2); Horse - EquCab3.0 (GCA_002863925.1); Dog - ROS_Cfam_1.0/Dog10K_Boxer_Tasha (GCA_014441545.1); Macaque - Mmul_10 (GCA_003339765.3); Cat - Felis_catus_9.0 (GCA_000181335.4); Pig - Sscrofa11.1 (GCA_000003025.6).

Approved genes are made public and searchable on our website https://vertebrate.genenames.org, which is updated on a daily basis. The full list of approved VGNC genes can be browsed and filtered by species and/or coding status (Figure 3). Information about each individual gene is displayed on “Symbol Report” pages, which include basic information about the gene, links to the corresponding NCBI and Ensembl gene annotations as well as links to specialist gene databases for that species if present, links to protein resources for the gene product, and links to named orthologs of the gene (Figure 3).

**Figure 3.**
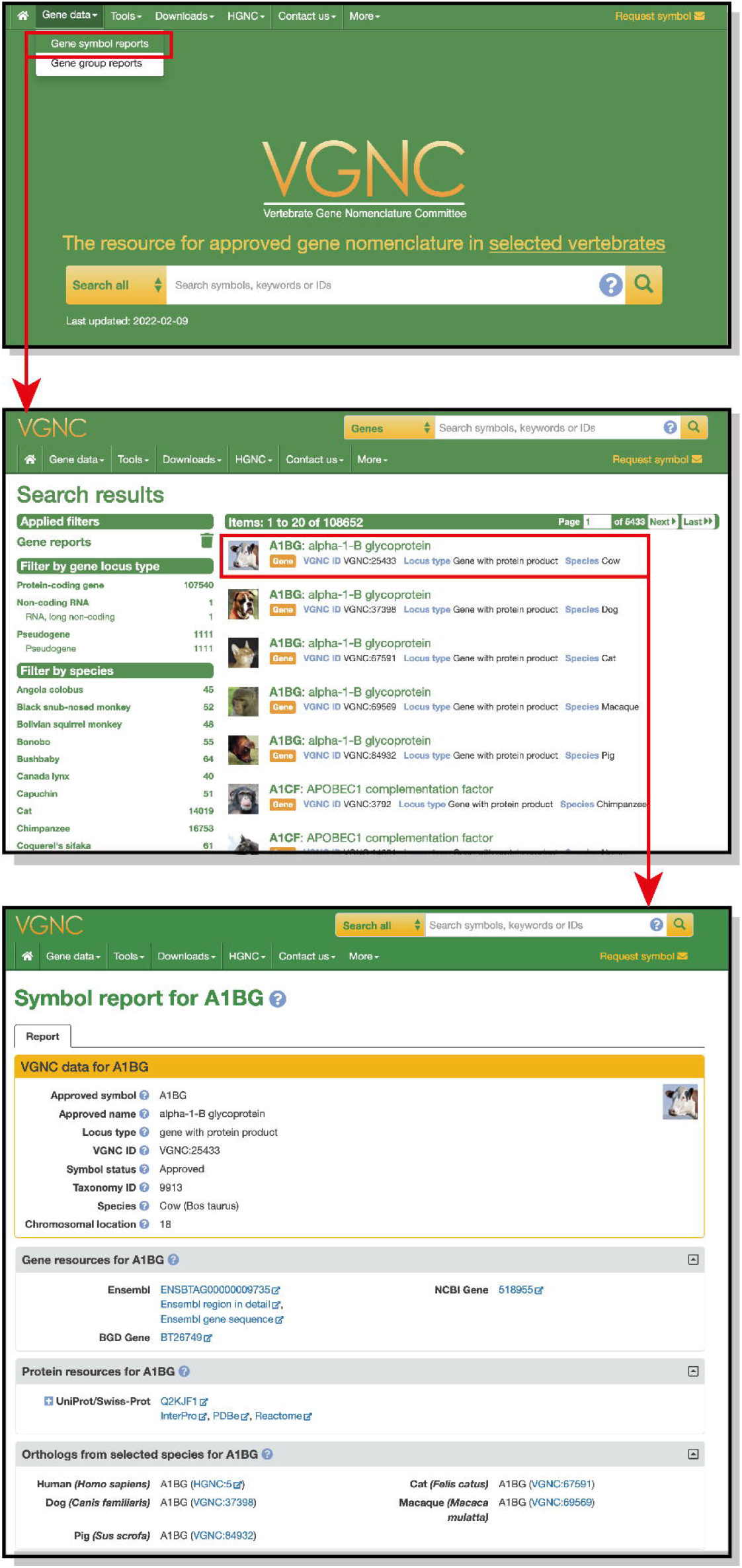
Screenshots showing the VGNC homepage, search result page and example gene symbol report. An example workflow is highlighted in red: Clicking on “Gene symbol reports” in the “Gene data” menu will run a search for all approved VGNC gene entries, shown in the second screenshot. Search results can be further filtered using the options on the left of the search results, and clicking on an individual result will take the user to the symbol report for that gene, shown in the third screenshot.

### Coverage of human genes with approved orthologs in VGNC

As of January 2022, there were 19,220 HGNC-approved protein-coding genes for human - 17,883 (93%) had at least 1 ortholog represented in the VGNC database but 1,367 did not yet have any ortholog approved in VGNC. We investigated each of these 1,367 genes to determine the reasons that they have not been approved in any VGNC species to date (Figure 4, Additional file 2: Table S2).

**Figure 4:**
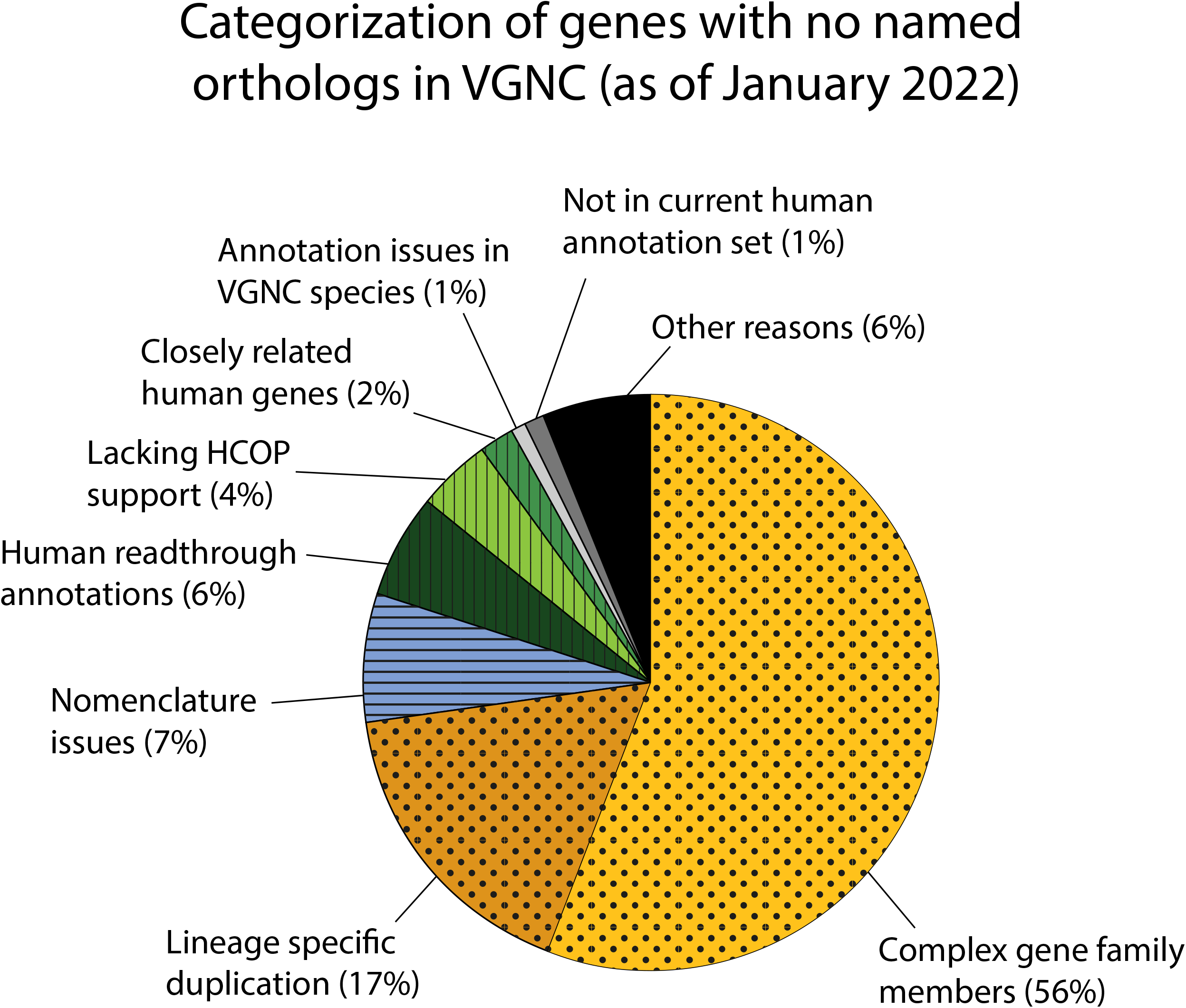
Pie chart categorizing reasons why there were no VGNC named orthologs for some human genes. As of January 2022 (N = 1,367). Complex gene family members = 759 genes; Lineage specific duplication = 234 genes; Nomenclature issues = 99 genes; Human readthrough annotations = 76 genes; Lacking HCOP Support = 54 genes; Closely related human genes = 34 genes; Annotation issues in VGNC species = 19 genes; Not in current human annotation set = 14 genes, Other = 78 genes. See article text for further explanation of each category and Additional file 2: Table S2 for full list of genes.

Broadly, the majority of these genes fell into the following categories: genes in large families or with copy number variation that required more detailed manual analysis before nomenclature assignment in vertebrate species (Fig. 4, yellow/dotted segments); genes for which the human nomenclature was unsuitable for transferral to other species (Fig. 4, blue/horizontal striped segments); and genes that likely have simple 1:1 orthology relationships across species but did not pass our automated orthology prediction threshold (3 out of 4 orthology assertions in Panther, NCBI Gene, Ensembl Compara and OMA) (Fig. 4, green/vertical striped segments).

In many cases the absence of a VGNC ortholog of a human gene is due to gene number variation causing uncertain orthology relationships, which is common in gene families that have frequent gene gains and losses throughout vertebrates. Large gene families often require manual curation including phylogenetic analysis of many genes across multiple species in order to assign nomenclature that accurately reflects evolutionary relationships. We found that 759 human protein coding genes do not yet have orthologs approved in VGNC due to their membership in a complex gene family (Fig. 4, “Complex gene family members”). Examples include genes encoding zinc finger containing proteins, keratins and interferons. Other examples that require manual input from a curator include genes that have undergone lineage specific duplication in humans or primates. We found that 234 human protein coding genes had not yet had a VGNC ortholog approved due to lineage specific duplications (Fig. 4, “Lineage specific duplication”). In these cases a decision must be made by a curator as to what nomenclature is most appropriate to assign to the non-duplicated orthologs.

The VGNC pilot project in which chimpanzee genes were manually approved provided an opportunity to review human gene nomenclature for suitability of use outside of humans. While it is preferable to use the same nomenclature for orthologous genes in different species to enable their quick identification, there are some human genes with nomenclature unsuitable for transfer to other species. This was recognised at the “Gene Nomenclature Across Species” meeting (9) where a key recommendation was that ‘humanizing’ nomenclature in other species should be avoided. Genes with human-centric nomenclature have been reviewed and the gene names updated while the gene symbol has been retained, where possible, often with the agreement of the communities working on them.

Examples include human disease specific gene names such as “malignant fibrous histiocytoma amplified sequence 1” (*MFHAS1*, HGNC:16982), which was renamed to “multifunctional ROCO family signaling regulator 1” to make it suitable for use across species, and names that included reference to other species, such as “dispatched homolog 1 (Drosophila)” (*DISP1*, HGNC:19711) which was renamed to “dispatched RND transporter family member 1”. There are still 99 genes with names referencing human disease that have not yet been renamed, and the orthologs of these genes have therefore not yet been approved in VGNC species (Fig. 4, “Nomenclature issues”).

We found that 54 human protein coding genes appear to have 1:1 orthology across VGNC species but did not pass our orthology prediction threshold for inclusion in the VGNC database, for reasons we could not identify (Fig. 4, “Lacking HCOP support”). These orthology relationships will require further investigation to confirm 1:1 orthology before approval in VGNC. In a further 76 cases, readthrough annotations on the human reference genome caused orthology prediction tools to fail to find 1:1 orthologs, since the non-human gene was predicted to have two human orthologs: the “true” ortholog and a readthrough annotation containing some or all of the same coding region (Fig. 4, “Human readthrough annotations”). All of these have since been manually reviewed and approved in at least one VGNC species. 19 human genes appear to have orthologs in VGNC species but problems with the gene annotations meant that they have not been automatically approved in the VGNC pipeline (Fig. 4, “Annotation issues in VGNC species”). 34 human genes appear to be single copy in all VGNC species but have closely related paralogs and so the orthology prediction resources could not distinguish between orthologs and paralogs across species (Fig. 4, “Closely related human genes”). 14 human genes that have been named by HGNC are not annotated on the current reference genome and so orthology prediction resources do not include these genes in their datasets (Fig. 4, “Not in current human annotation set”).

Further, less common, reasons for a gene having no VGNC ortholog approved were combined into a final category of “Other” (Fig. 4). This includes genes for which there is no consensus on the locus type in humans between Ensembl, NCBI and HGNC, i.e. it is unknown whether the gene is protein-coding or not. The “Other” category also includes a small number of genes in complex immune-related families where 1:1 orthologs do not exist across species, such as the killer immunoglobulin-like receptors and major histocompatibility complex genes. Although other species have members of these gene families, there is no 1:1 orthology and so unique gene symbols will be approved in each species (17).

### Naming pseudogenes in VGNC

The VGNC has not prioritized the systematic naming of pseudogenes across multiple species, however there are some specific examples of pseudogenes receiving approved nomenclature: large gene families such as the olfactory receptors, cytochrome P450s and histones have a significant proportion of pseudogenes, and any pseudogenes within these families with NCBI/Ensembl annotations, that have also been manually curated by our expert collaborators or VGNC curators, have been named. The VGNC database also includes some pseudogenes that were initially approved as protein coding but their gene models have since been updated to pseudogenes.

An area of particular interest for the VGNC has been approving orthologs of genes that are pseudogenized in humans but coding in other species (so-called “unitary” pseudogenes). These are genes that would otherwise not receive approved nomenclature via automated means, because vertebrate gene naming is often based on the human ortholog and orthology prediction algorithms do not typically include pseudogenes in their data sets, and therefore vertebrate orthologs need to be manually identified. There are currently 274 HGNC approved human pseudogenes that have protein coding orthologs in other species and have been named as such. The majority of these pseudogenes were initially named relative to mouse protein coding orthologs. To date we have approved nomenclature for multiple orthologs of 104 human unitary pseudogenes (Additional file 3: Table S3) and will continue to prioritize these genes in our manual curation.

### Gene groups in VGNC

The VGNC recently introduced a feature called “Gene Groups” which has been a part of the HGNC database for over 20 years. Human genes are grouped based on shared characteristics such as homology, structure, common functions and/or phenotypes, and protein complex membership. We have introduced a subset of these Gene Groups to VGNC where we have completed considerable VGNC curation of large gene families, ie, the Olfactory Receptors, Keratins, Histones and Cytochrome P450s. Gene Group reports allow visualization and navigation of hierarchical groups in complex gene families (Figure 5).

**Figure 5.**
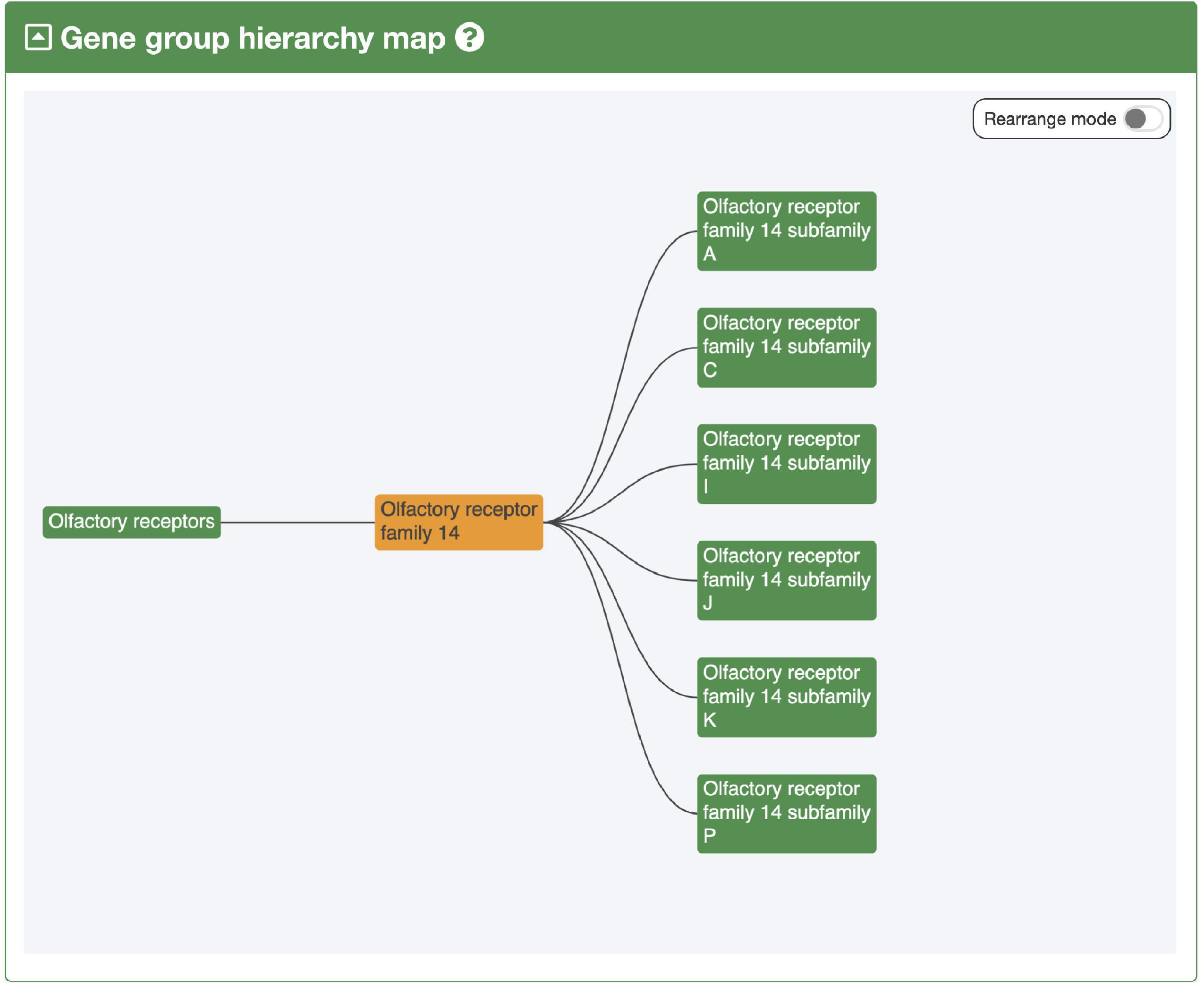
Example of the VGNC gene group hierarchy navigator for Olfactory receptor family 14. Gene group hierarchies are displayed on gene group pages and show the parent and child groups of the gene group of interest. In this example, we can see that Olfactory receptor family 14 has the parent group “Olfactory receptors” and 6 child groups representing its subfamilies. The groups shown in the hierarchy diagram are clickable and so can be used to navigate through a hierarchical gene group.

### VGNC data dissemination

VGNC approved nomenclature is automatically imported and displayed by the NCBI (12), Ensembl (11) and UniProt (18) databases. This ensures that once a gene has been approved in VGNC it has a consistent gene symbol and name across these resources and is visible to the community even if they are not VGNC website users. An additional benefit of this nomenclature dissemination is that as more accurate nomenclature is included for key vertebrate species in NCBI and Ensembl, the more likely it is that the appropriate nomenclature will be assigned in these databases’ automated nomenclature pipelines even for non-VGNC curated species, as these pipelines are based on orthology between species.

## Discussion and Future Plans

The VGNC has approved nomenclature for over 100,000 genes in a variety of vertebrate species, and the approved nomenclature is disseminated widely via major biological databases. The benefits of approved nomenclature extend beyond just the species included in the VGNC, as both NCBI and Ensembl use homology to automatically project nomenclature to other species in their databases. The VGNC project has also led to large scale revision of nomenclature for complex gene families, for example, completely independent naming systems were in place for the olfactory receptor genes in different species (15), which made it impossible to determine orthology and paralogy relationships based on nomenclature. Our ongoing efforts to harmonize olfactory receptor gene nomenclature across species will make homology relationships obvious at a glance.

VGNC manual curation has resulted in improvements to gene nomenclature that would not have been possible using current automated techniques. Several species-specific duplications of genes or regions have been identified and assigned novel nomenclature: for example, there has been a tandem duplication in the human lineage leading to duplication of the *MMP23* gene and subsequent pseudogenization of one of the copies. The intact human gene has the symbol *MMP23B* and the pseudogene has the symbol *MMP23AP*. Since orthology prediction algorithms typically only include coding sequences, the single copy orthologs in other species were being automatically assigned identical nomenclature to human *MMP23B* in other resources. VGNC manual curation allowed identification of this issue and thus the non-human orthologs have now been correctly approved as *MMP23*. Similarly, VGNC has now manually approved nomenclature for at least 104 orthologs of human pseudogenes that would not have received approved nomenclature by other means (Additional file 3: Table S3).

Manual curation has also led to corrections in data beyond gene nomenclature. It has been possible to identify issues with automatically predicted gene models in NCBI and Ensembl annotation sets such as merging of neighboring genes or fragmented gene models. Correspondence with RefSeq curators at NCBI has allowed for periodic review and correction of gene models in their database which then allows VGNC nomenclature to be approved and linked to a corrected NCBI Gene ID.

Challenges faced in the VGNC project include the use of different genome assembly versions between NCBI and Ensembl, making it more difficult for curators to compare gene models and synteny in the two different versions since coordinates and annotations differ across assemblies. For example, NCBI and Ensembl have been annotating different versions of the chimpanzee genome since 2018. We currently do not approve gene nomenclature if we cannot link to a suitable gene model in either NCBI or Ensembl, and at present there is no provision within Ensembl to make routine manual corrections to gene models in species outside of human and mouse.

A large majority of the genes approved in VGNC are those in which orthology has been easily determined using the approach described and thus have been able to be automatically approved or quickly manually approved. Manual curation is time consuming and hence our approach so far has been to concentrate our efforts to maximise the number of genes approved. More recently we have focused on complex gene families, taking a multidisciplinary approach to assign nomenclature across multiple species. This often occurs in collaboration with other nomenclature authorities such as the MGNC, RGNC, CGNC, Xenbase and ZNC as previously described. These efforts will be coupled with expansion to the Gene Groups feature in VGNC, including the addition of both more Gene Groups and their members.

The VGNC’s remit to date has been to assign nomenclature to coding genes, but in future we intend to explore the naming of non-coding genes, including pseudogenes and non-coding RNA genes. This will likely be limited to non-coding genes that are either highly conserved across species or have been characterized in the literature. Nomenclature approval for non-coding RNA genes will begin with microRNA genes. Human microRNA genes are currently assigned gene symbols as a result of a long standing collaboration between the HGNC and miRBase (19). MicroRNA identifiers are provided by miRBase and follow the format mir-# (e.g. mir-17) for the stem loop and miR-# (e.g. miR-17) for the mature miRNA, while HGNC approves the format MIR# for the encoding gene (e.g. *MIR17*). Equivalent gene symbols are currently approved for mouse and rat microRNA genes by MGNC and RGD; the mouse and rat orthologs of human *MIR17* have the gene symbol *Mir17*. In future, we will look into incorporating microRNA orthology predictions to approve symbols for microRNA genes that are orthologous to human microRNA genes for our seven core species. We will also explore approving symbols for long non-coding RNA (lncRNA) genes that have HGNC-curated mouse orthologs and have been approved unique symbols from publications. This would be a small number of lncRNA genes and we would not expect that adequately annotated orthologs would be present in all key species to allow approval of VGNC symbols.

Other future improvements we have planned for the VGNC include the development of tools to improve curation efficiency, for example, based on synteny across multiple species. We also plan to implement additional quality control tools to allow curators to quickly identify data changes that affect approved gene nomenclature, for example, when gene annotation identifiers are changed in NCBI and Ensembl.

## Conclusions

As the number of vertebrate genomes continues to increase with large scale sequencing projects such as the Vertebrate Genomes Project (20)), this provides a unique opportunity to assign gene nomenclature across species that reflects the evolutionary history of genes and gene families, while remaining consistent with existing human gene nomenclature used in clinical settings. It is important that gene nomenclature assignment is carried out in a coordinated manner with the appropriate gene nomenclature authorities to ensure that genes are labelled in a way that facilitates unambiguous communication, and we strongly encourage researchers to work with the gene nomenclature authorities such as VGNC when proposing novel gene nomenclature in vertebrate species.

## Declarations

### Ethics approval and consent to participate

Not applicable

### Consent for publication

Not applicable

### Availability of data and materials

All data generated or analysed during this study are either publicly available at https://vertebrate.genenames.org, or included in this published article and Additional files.

### Authors’ Contributions

EAB conceived the project, BY and KG designed and maintained the database schema and website, TEMJ, BB, EAB, RLS and ST analysed data, TEMJ wrote the first draft and all authors contributed to the writing of the manuscript.

## Acknowledgements

The VGNC would like to thank past and present members of the HGNC and VGNC group, especially Paul Denny for his input on histone gene naming. We are grateful for the contributions of expert collaborators who provided data for specific gene families: David Nelson, Tsviya Olender and Doron Lancet. Finally we thank the HGNC/VGNC Scientific Advisory Board members past and present who have offered their valuable input into the project’s implementation.

## Additional Files

Additional file 1.xlsx : Table S1: Summary of cytochrome P450 genes approved in VGNC.

Additional file 2.xlsx : Table S2: HGNC genes without VGNC approved orthologs as of January 2022.

Additional file 3.xlsx : Table S3: VGNC approved orthologs of human unitary pseudogenes.

